# Negeviruses reduce replication of alphaviruses during co-infection

**DOI:** 10.1101/2020.09.01.277517

**Authors:** Edward I. Patterson, Tiffany F. Kautz, Maria A. Contreras-Gutierrez, Hilda Guzman, Robert B. Tesh, Grant L. Hughes, Naomi L. Forrester

## Abstract

Negeviruses are a group of insect-specific virus (ISV) that have been found in many arthropods. Their presence in important vector species led us to examine their interactions with arboviruses during co-infections. Wild-type negeviruses reduced the replication of Venezuelan equine encephalitis virus (VEEV) and chikungunya virus (CHIKV) during co-infections in mosquito cells. Negev virus (NEGV) isolates were also used to express GFP and anti-CHIKV antibody fragments during co-infections with CHIKV. NEGV expressing anti-CHIKV antibody fragments was able to further reduce replication of CHIKV during co-infections, while reductions of CHIKV with NEGV expressing GFP were similar to titers with wild-type NEGV alone. These results are the first to show that negeviruses induce superinfection exclusion of arboviruses, and to demonstrate a novel approach to deliver anti-viral antibody fragments with paratransgenic ISVs. The ability to inhibit arbovirus replication and express exogenous proteins in mosquito cells make negeviruses a promising platform for control of arthropod-borne pathogens.

## Introduction

Many insect-specific viruses (ISVs) have been discovered in wild-caught and laboratory colonies of mosquitoes and in mosquito cell cultures [1]. ISVs are only known to replicate in arthropods or insect cell lines. While posing no threat to human or animal health, ISVs may affect the transmission of more dangerous vector-borne pathogens. Highly insect-pathogenic ISVs have been suggested for use as biological control agents to reduce populations of vector competent mosquitoes [2-4]. Several recent studies have demonstrated that ISVs may play a more direct role by inhibiting the replication of arboviruses within the insect host. The majority of these experiments have attempted to define a relationship based on superinfection exclusion, a phenomenon in which an established virus infection interferes with a secondary infection by a closely related virus. For example, insect-specific flaviviruses, such as cell fusing agent virus (CFAV), Nhumirim virus (NHUV) and Palm Creek virus (PCV) have demonstrated an ability to reduce viral loads of vertebrate pathogenic flaviviruses, like West Nile virus (WNV), Zika virus (ZIKV), dengue virus (DENV), Japanese (JEV) and St. Louis encephalitis (SLEV) viruses [5-10]. Similarly, the insect-specific alphavirus Eilat virus (EILV) was shown to reduce or slow replication of the pathogenic alphaviruses chikungunya virus (CHIKV), Sindbis virus (SINV), eastern (EEEV), western (WEEV) and Venezuelan equine encephalitis (VEEV) viruses in cell culture or in mosquitoes [11]. Less information is available about the effect of unrelated viruses during superinfection. Cell cultures chronically infected with *Aedes albopictus* densovirus (AalDNV) limit replication of DENV [12], and cell cultures with established CFAV and Phasi Charoen-like virus (PCLV) infections reduced ZIKV and DENV replication [13]. The mechanism for these reduced titers has not been elucidated, but the relationships appear to be virus specific and even host specific [5, 14, 15].

The genus *Negevirus* is a recently discovered, unclassified group of ISVs [16]. Members of this genus have been isolated from several species of hematophagous mosquitoes and sandflies, and negev-like viruses have also been found in other non-vector insects [17-24]. Phylogenetic studies have placed this group of viruses most closely to members of the genus *Cilevirus*, plant pathogens that are transmitted by mites [16, 23]. These viruses have a single-stranded, positive-sense RNA genome of ∼9-10 kb, and contain three open reading frames (ORFs) [16]. The ORFs encode for the replication machinery (ORF1), a putative glycoprotein (ORF2), and a putative membrane protein (ORF3). Electron microscopy has shown the structural proteins to be arranged in a hot air balloon morphology, a round particle with a single protrusion that is likely the glycoprotein structure [25-27]. Little is known about the infectivity, transmission dynamics, and species range of negeviruses. However, they are commonly found in field collected mosquitoes [28, 29].

The association of negeviruses with important vector species over a wide geographical range raises the question of possible interactions or interference of negeviruses with vertebrate pathogenic viruses. Few studies exist that demonstrate the ability of unrelated viruses to induce superinfection exclusion, but evidence for this phenomenon with negeviruses could provide a platform to control vector-borne viral diseases in many arthropod vector species. In this study, three negevirus isolates from the Americas were assessed for superinfection exclusion in cell cultures with CHIKV and VEEV. The use of a Negev virus (NEGV) infectious clone also allowed manipulation of the virus genome to provide a greater ability to exclude superinfection with these pathogenic alphaviruses.

## Materials and Methods

### Cell culture and viruses

*Aedes albopictus* (C7/10) cells [30] were obtained from the World Reference Center for Emerging Viruses and Arboviruses (WRCEVA). African green monkey kidney (Vero E6) cells were obtained from the American Type Culture Collection (ATCC). C7/10 cells were maintained in Dulbecco’s minimal essential medium (DMEM) supplemented with 10% fetal bovine serum (FBS), 1% minimal essential medium non-essential amino acids, 1% tryptose phosphate broth and 0.05 mg/mL gentamycin in a 30°C incubator with 5% CO_2_. Vero cells were maintained in DMEM supplemented with 10% FBS and 0.05 mg/mL gentamycin in a 37°C incubator with 5% CO_2_.

Negev virus (NEGV) was rescued in C7/10 cells from an infectious clone as previously described and without further passage [31]. The sequence was derived from NEGV strain M30957 isolated from a pool of *Culex coronator* mosquitoes collected in Harris County, Texas, USA in 2008 [16]. Piura virus (PIUV) strain EVG 7-47 (PIUV-Culex) isolated from a pool of *Culex nigripalpus* mosquitoes from Everglades National Park, Florida, USA in 2013 [21]. PIUV EVG 7-47 was passaged four times in C6/36 cells and obtained from the WRCEVA. PIUV strain CO R 10 (PIUV-Lutzomyia) was isolated from a pool of *Lutzomyia evansi* sandflies caught in Ovejas, Sucre, Colombia in 2013 [21]. The isolate PIUV CO R 10 was passaged twice in C6/36 cells and also obtained from the WRCEVA. CHIKV isolate 181/25 [32] was rescued in Vero cells from an infectious clone as previously described [33]. Rescued CHIKV 181/25 was subsequently passaged once in C7/10 cells and once in Vero E6 cells. Venezuelan equine encephalitis virus (VEEV) vaccine strain, TC-83 [34], was rescued in baby hamster kidney (BHK) cells from an infectious clone without further passage.

### Cloning NEGV for exogenous gene expression

The NEGV infectious clone was used as the backbone to express exogenous genes. Green fluorescent protein (GFP) or the single chain variable fragment (scFv) of anti-CHIKV neutralizing antibody CHK265 [35] was inserted on the C-terminal of ORF3 as either a fusion protein, or with a 2A sequence (EGRGSLLTCGDVEENPGP) (Figure 1A). The cloned scFv CHK265 sequence contained a N-terminal linker (LAAQPAMA) for articulation from the viral ORF3 protein, and a domain linker ((G_4_S)_4_) between the variable heavy (V_H_) and variable light (V_L_) domains (Integrative DNA Technologies) (Figure 1B). Cloning was performed using In-Fusion HD Cloning Kit (Takara Bio) as per the manufacturer’s protocol. Correct insertion was confirmed by sequencing. Infectious clones of NEGV containing exogenous genes were rescued in C7/10 cells as previously described and without further passage [31].

**Figure 1.**
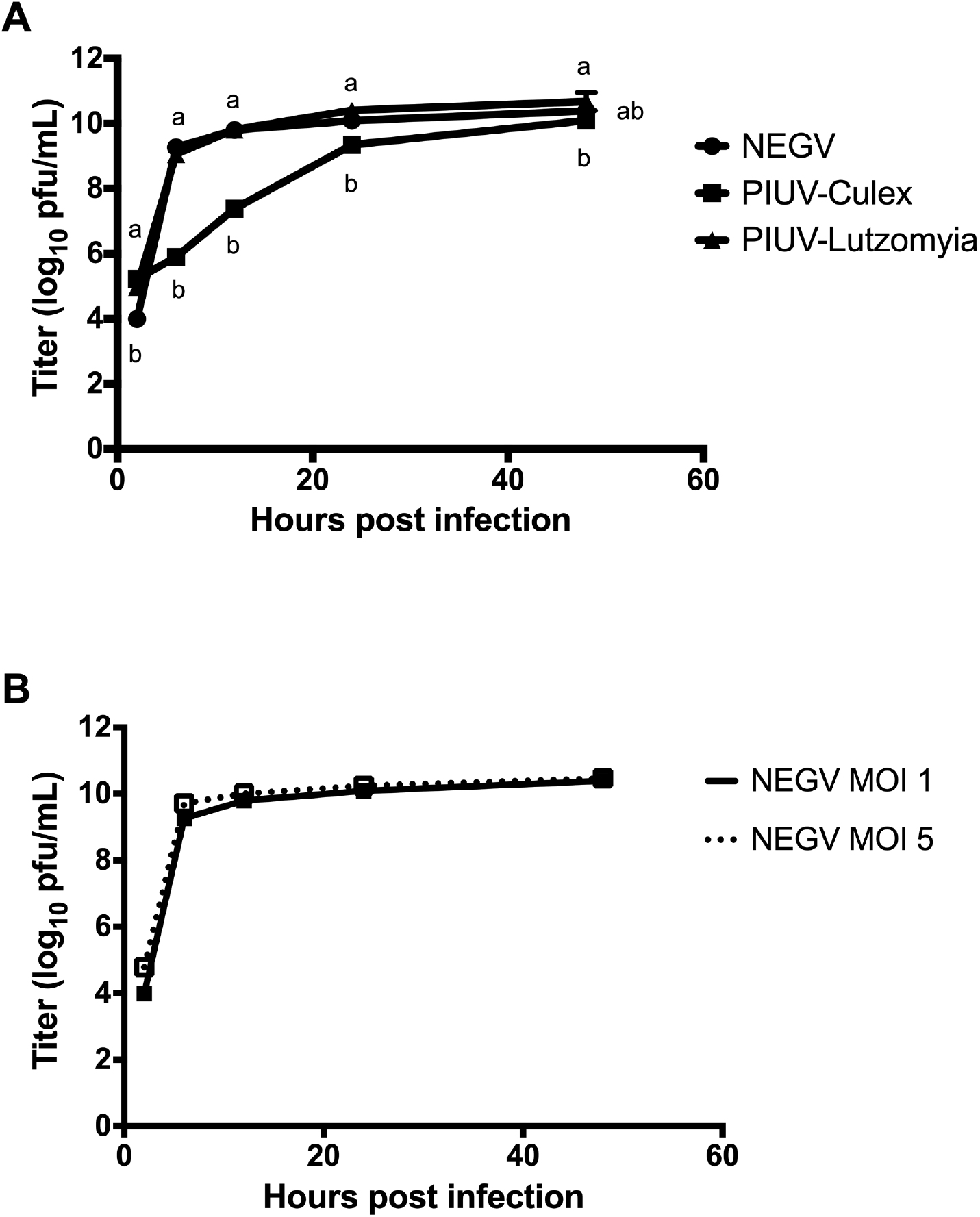
Growth curve for wild-type negeviruses. A) The titer of each virus, Negev virus (NEGV), Piura virus-Culex (PIUV-Culex) and Piura virus-Lutzomyia (PIUV-Lutzomyia) at different time points following infection at MOI of 1 in C7/10 cells. B) Growth curve of NEGV with MOI of 1 and 5 in C7/10 cells. All points represent mean of n=3, ± SD. Letters indicate significant differences (p < 0.0001).

### Virus growth curves

Negevirus and alphavirus growth curves were done in C7/10 cells maintained at 30°C and 5% CO_2_. Negeviruses were inoculated at a multiplicity of infection (MOI) of 1 or 5. Alphaviruses were inoculated at a MOI of 0.1. Virus was added to the cells which were incubated at 30°C for one hour. Inoculum was removed, cells were washed with PBS, and fresh media was added to the wells. Cells were incubated in a 30°C incubator with 5% CO_2_. Samples were collected in triplicate at 2-, 6-, 12-, 24- and 48-hours post infection (hpi). Samples were clarified by centrifugation at 1962 *x g* for 5 minutes. Supernatant was removed and stored at −80°C until used for plaque assays. Negevirus titers were determined by plaque assay in C7/10 cells as previously described [31]. Alphavirus titers were determined by standard plaque assay in Vero E6 cells.

### Negevirus-alphavirus co-infections

C7/10 cells were inoculated with negevirus isolates at a MOI of 1 or 5. The cells were also inoculated with an alphavirus at a MOI of 0.1 at 0, 2, or 6 hours post negevirus infection. Media was removed after 1 hour of simultaneous incubation with negevirus and alphavirus inocula. Cells were then washed with PBS, and fresh media was added to the wells. Cells were held in a 30°C incubator with 5% CO_2_. Samples were collected in triplicate at 12-, 24- and 48-hours post alphavirus infection. Samples were clarified by centrifugation at 1962 *x g* for 5 minutes. Supernatant was removed and stored at −80°C until used for plaque assays. Alphavirus titers were determined by standard plaque assay in Vero E6 cells.

### Statistical analysis

Differences in virus growth curves were determined by two-way ANOVA followed by Tukey’s test. Comparison of NEGV growth curves with different MOIs was determined by multiple t tests followed by Holm-Sidak method. All statistical tests were performed using GraphPad Prism 6.0.

## Results

### Wild-type negevirus growth curves

All wild-type negeviruses reached titers greater than 10log_10_ pfu/mL within 48 hours when infected at a MOI of 1 (Figure 1A). NEGV and PIUV-Lutzomyia neared peak titer by 12 hours post infection (hpi), while PIUV-Culex neared peak titer at 24 hpi. Infections of NEGV with MOIs of 1 and 5 produced similar growth curves (Figure 1B).

### Superinfection exclusion of alphaviruses with wild-type negeviruses

To determine the effect of negeviruses on the replication of alphaviruses in cell culture, negevirus isolates were co-infected with VEEV or CHIKV isolates. NEGV was able to significantly reduce replication of VEEV, with reductions of 5.5-7.0log_10_ pfu/mL of VEEV at 48 hours (Figure 2A). The MOI of NEGV and timing of VEEV inoculation had no significant difference on the reduction. Co-infection with PIUV-Culex or PIUV-Lutzomyia also significantly reduced replication of VEEV across all time points (Figure 2B-D). A similar reduction of VEEV was observed during all negevirus co-infections, as VEEV was reduced 4.6-7.2log_10_ pfu/mL at 48 hours.

**Figure 2.**
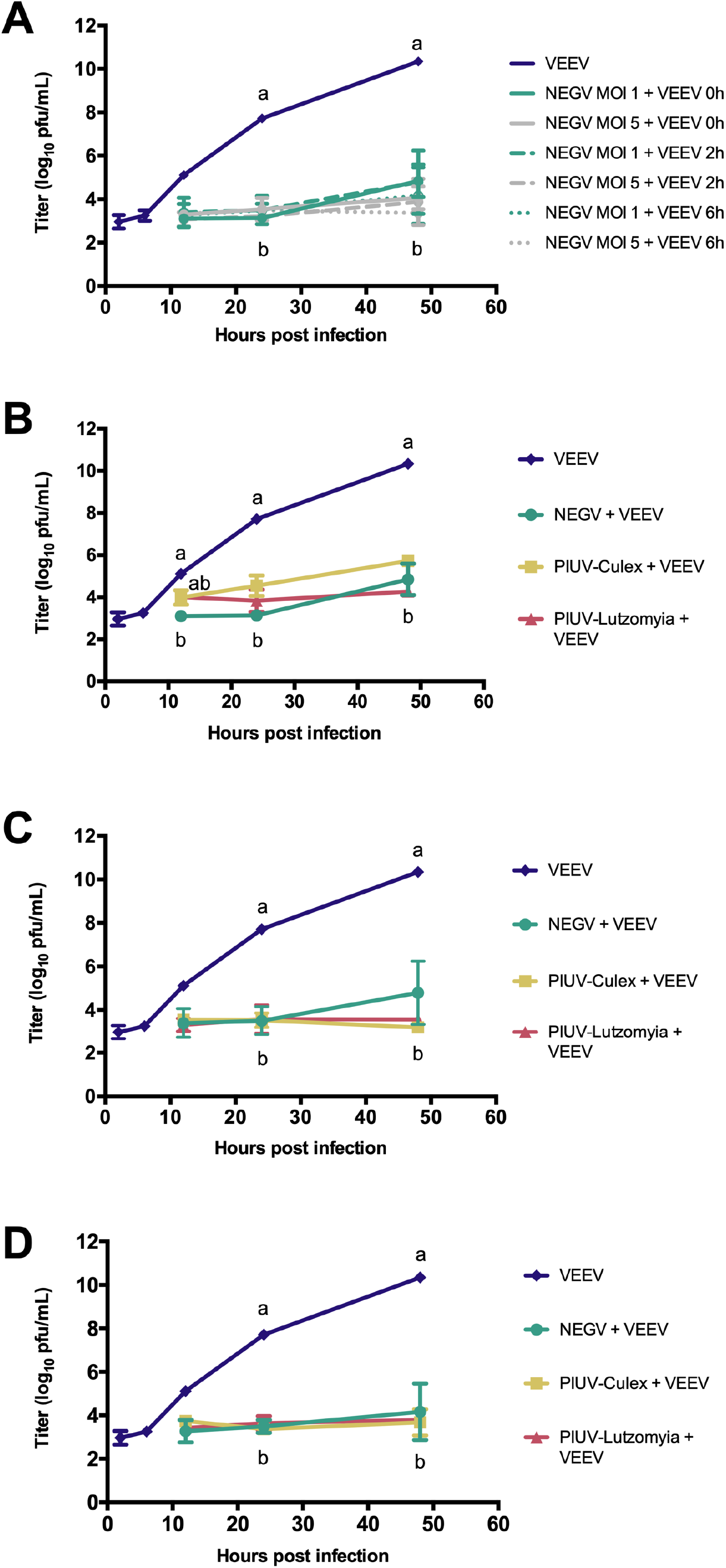
Growth curves of VEEV in C7/10 cells during co-infections with wild-type negeviruses. A) Growth curves of VEEV when inoculated on cells at 0-, 2- and 6 hours post NEGV infections. NEGV was inoculated at MOI 1 or 5. B) Growth curves of VEEV when inoculated on cells at 0 hours post negevirus infection, C) 2 hours post negevirus infection, and D) 6 hours post negevirus infection. Negeviruses were inoculated at MOI of 1. All points represent mean of n=3, ± SD. Letters indicate significant differences (p < 0.0001).

Co-infections with CHIKV and NEGV also resulted in significantly lower titers of CHIKV at all time points, but only reducing the titer of CHIKV by 0.65-0.93log_10_ pfu/mL after 48 hours (Figure 3A). Varying the MOI of NEGV and timing of CHIKV inoculation only produced differing titers of CHIKV at the 12-hour timepoint. However, titers of CHIKV during co-infection with different negeviruses varied greatly (Figure 3B, C), with the largest variance of CHIKV titers, reductions of 0.7log_10_, 2.4log_10_ and 5.3log_10_ pfu/mL, observed when inoculated 6 hours post-inoculation with NEGV, -PIUV-Culex and -PIUV-Lutzomyia, respectively (Figure 3D).

**Figure 3.**
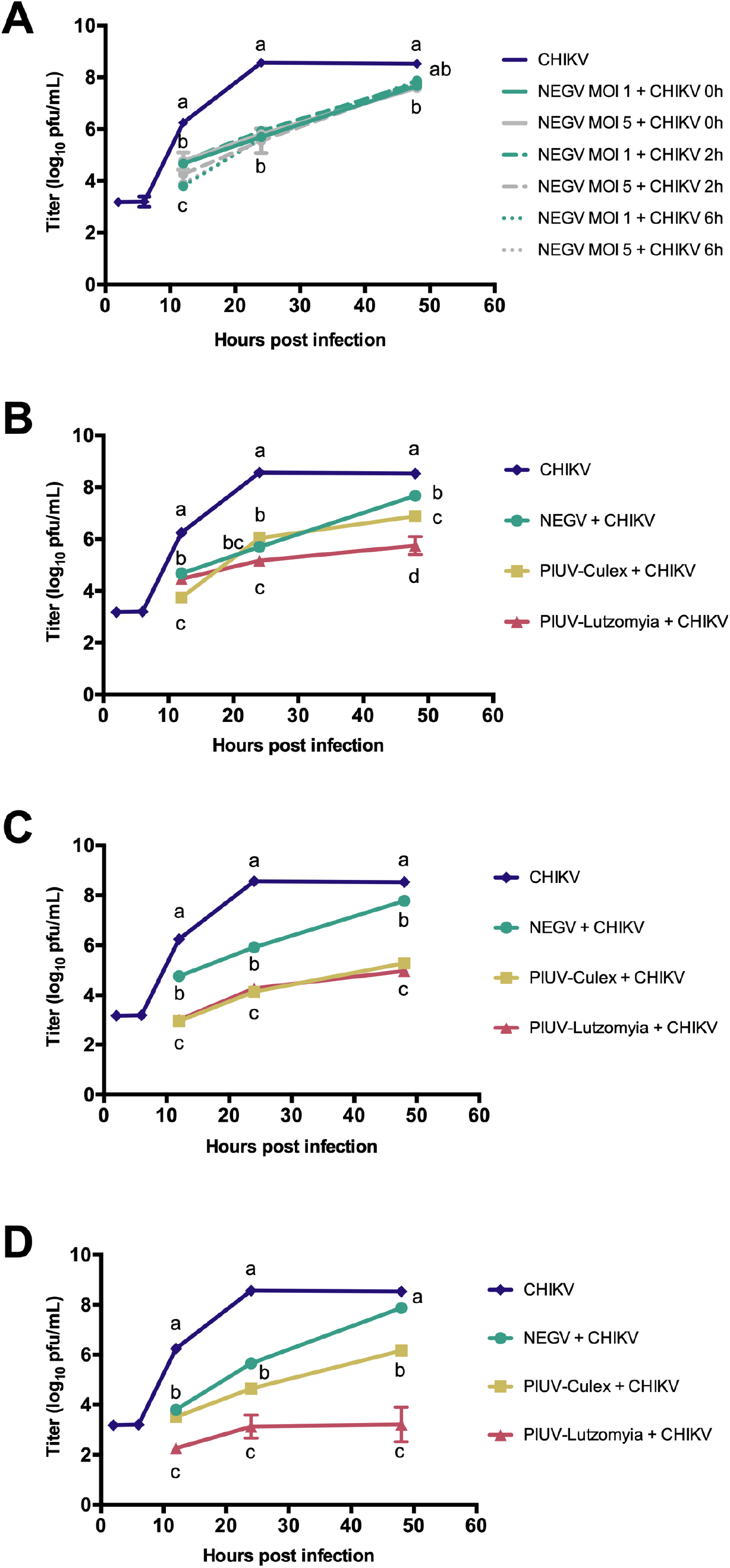
Growth curves of CHIKV in C7/10 cells during co-infections with wild-type negeviruses. A) Growth curves of CHIKV when inoculated on cells at 0-, 2- and 6 hours post NEGV infections. NEGV was inoculated at MOI 1 or 5. B) Growth curves of CHIKV when inoculated on cells at 0 hours post negevirus infection, C) 2 hours post negevirus infection, and D) 6 hours post negevirus infection. Negeviruses were inoculated at MOI of 1. All points represent mean of n=3, ± SD. Letters indicate significant differences (p < 0.0001).

### Replication of modified NEGV isolates

Sequences for GFP and scFv CHK265, an anti-CHIKV antibody, were cloned as both fusion-and cleaved proteins on ORF3 of the NEGV infectious clone (Figure 4A). Mutated isolates were rescued and had titers ranging from 9.6-10.4log_10_ pfu/mL, with similar growth curves to the wild-type NEGV (Figure 4B). Cells infected with NEGV GFP-fusion demonstrated brilliant, punctate fluorescence (Figure 4C), while cells infected with NEGV GFP (cleaved) demonstrated dull, diffuse fluorescence (Figure 4D).

**Figure 4.**
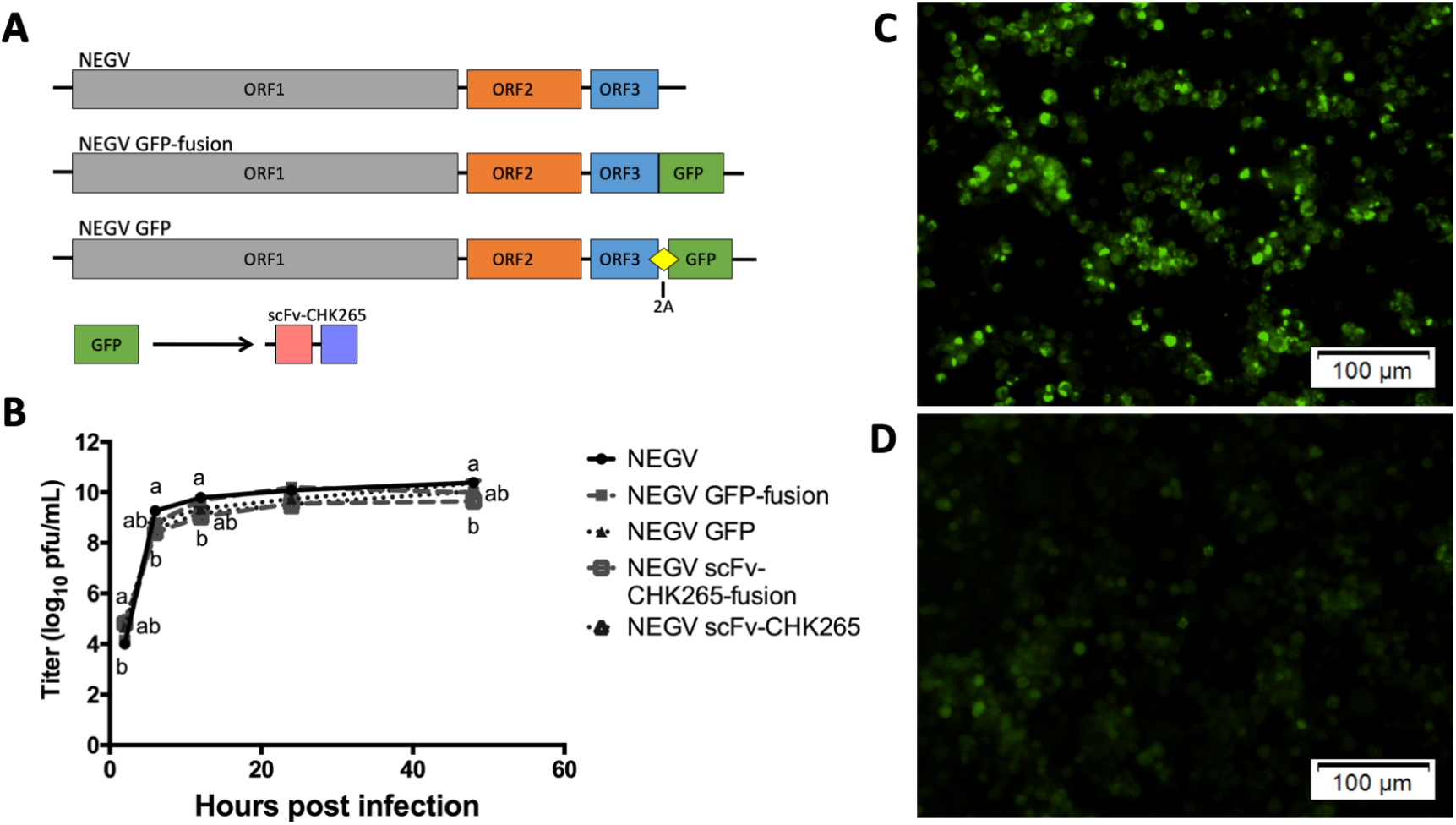
Rescued paratransgenic NEGV infectious clones. A) Schematic of NEGV genomes for wild-type, and GFP-expressing viruses. NEGV GFP-fusion added the GFP sequence onto ORF3 and NEGV GFP separated the ORF3 and GFP with a 2A sequence to produce the proteins separately. GFP was replaced by scFv-CHK265 for NEGV scFv-CHK265-fusion and NEGV scFv-CHK265. B) Growth curves of NEGV wild-type and NEGV mutants expressing GFP or scFv-CHK265. All points represent mean of n=3, ± SD. Letters indicate significant differences (p < 0.0001). C) Fluorescent microscopy of C7/10 cells infected with NEGV GFP-fusion. Cells demonstrate brilliant, punctate fluorescence. D) Fluorescent microscopy of C7/10 cells infected with NEGV GFP. Cells demonstrate dull, diffuse fluorescence.

### Superinfection exclusion of alphaviruses with modified NEGV

NEGV isolates expressing GFP or scFv-CHK265 were used to infect cells for co-infection with CHIKV. When infected simultaneously, CHIKV titers were reduced by 0.7-1.1 log_10_ pfu/mL during co-infections of NEGV expressing GFP, and by 2.9-3.8log_10_ pfu/mL during co-infections of NEGV expressing scFv-CHK265 at the 48-hour timepoint (Figure 5A). When inoculated 2 hours post NEGV infection, the titer of CHIKV after 48 hours was reduced 0.7-0.9log_10_ pfu/mL with NEGV expressing GFP and 3.7-4.5log_10_ pfu/mL with NEGV expressing scFv-CHK265 (Figure 5B). Delaying CHIKV infection 6 hours post NEGV resulted in reductions of 1.2-1.9log_10_ pfu/mL and 5.2-5.7log_10_ pfu/mL after 48 hours of co-infection with NEGV expressing GFP and scFv-CHK265, respectively (Figure 5C).

**Figure 5.**
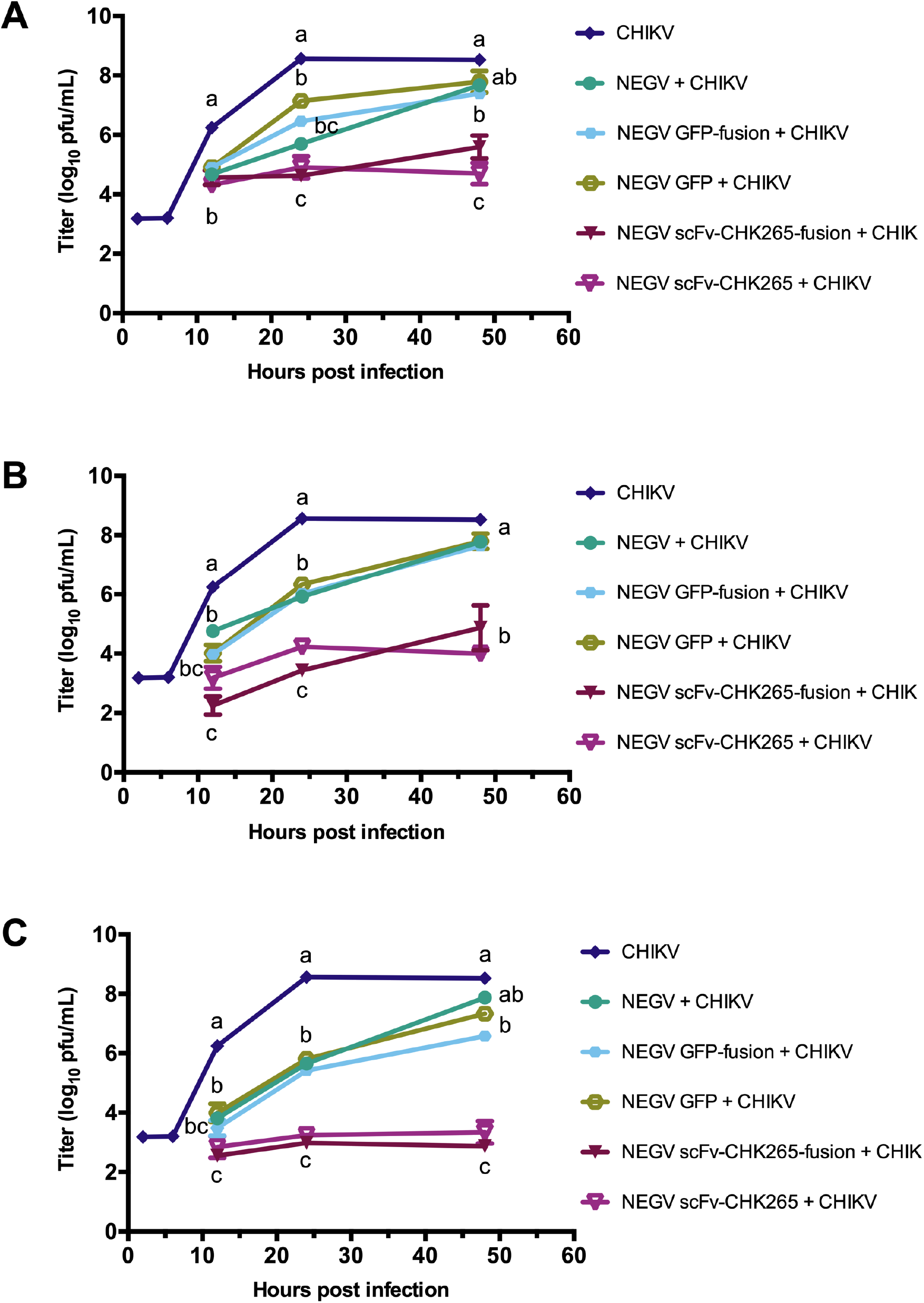
Growth curves of CHIKV during co-infections with paratransgenic NEGV. Growth curves of CHIKV when inoculated on cells at A) 0 hours post NEGV infection, C) 2 hours post NEGV infection, and D) 6 hours post NEGV infection. All NEGV isolates were inoculated at MOI of 1. All points represent mean of n=3, ± SD. Letters indicate significant differences (p < 0.0001).

## Discussion

The microbiome of arthropod vectors is known to influence host-pathogen interactions [36-38]. The precise mechanisms of pathogen inhibition are unknown, but there is increasing evidence that interference from ISVs is one mechanism [5-7, 10]. Interactions between related viruses has led to the theory of superinfection exclusion, in which an established infection interferes with or inhibits a secondary infection by a closely related virus. For example, a CFAV mosquito isolate reduced the replication of DENV and ZIKV during co-infections in mosquitoes and mosquito cells [5].

To investigate if superinfection exclusion occurred with other virus combinations, two pathogenic alphaviruses, VEEV and CHIKV, and three negeviruses, NEGV, PIUV-Culex and PIUV-Lutzomyia, were used in co-infection experiments. While VEEV titers were consistently reduced during all co-infection experiments with negeviruses, reductions varied during CHIKV-negevirus co-infections. These results provide further evidence that superinfection exclusion of alphaviruses is pathogen specific. Previous reports demonstrated no reduction in titer of VEEV TC-83 but significant reductions of wild-type VEEV and CHIKV after 48 hours when co-infected with EILV, an alphavirus ISV [11]. However, the potential for superinfection exclusion of pathogens is different for each ISV, despite their relatedness. These differences have been demonstrated among several insect-specific flaviviruses, Nhumirim virus (NHUV) and Palm Creek virus (PCV) were capable of superinfection exclusion; CFAV gave varying results; and Culex flavivirus (CxFV) did not reduce titers of pathogenic arboviruses [5-10, 14, 15, 39-42]. In our experiments, only some of the negeviruses tested were capable of inhibiting important arboviruses.

While ISVs show promising results to block arbovirus replication in mosquito vectors, their unknown mechanism of action may limit their use against a wide range of pathogens, but paratransgenic ISVs could be used to provide antiviral molecules that specifically interfere with pathogen transmission [43]. To this end, we used an infectious clone of NEGV to deliver a fragment of an antibody known to neutralize CHIKV. An scFv consists of the variable regions of the heavy and light chains of an antibody, joined by a soluble linker. These antibody fragments can possess the neutralizing qualities of their full-size versions in only ∼27kDa. Co-infections with scFv-expressing NEGV isolates greatly reduced titers of CHIKV, whereas co-infections with control NEGV isolates expressing GFP or wild-type NEGV only modestly reduced CHIKV titers. The use of parastransgenic NEGV expressing scFvs demonstrates a novel approach to disrupt pathogen infection in mosquitoes. This method adapts two existing techniques for pathogen control: *Wolbachia* infected mosquitoes and the CRISPR-Cas-aided integration of scFv sequences into the mosquito genome. *Wolbachia* is a ubiquitous species of bacteria found in many insects that has been shown to block replication of some viral pathogens in cell cultures and mosquitoes. The use of *Wolbachia*-infected vectors has been widely adapted to curb mosquito-borne viral diseases, propelled by its natural ability to colonize mosquitoes [44]. Negeviruses also possess this attribute, having been discovered in numerous mosquito species on 6 continents, along with sandflies and other diverse insect species [19-22, 24]. Insertion of gene-editing scFv sequences into mosquito genomes has also been used to prevent *Plasmodium* and DENV infection [45, 46]. By using CRISPR-Cas9 to insert a scFv targeting *Plasmodium*, infection was blocked in *Anopheles* mosquitoes, and gene drive insured the production of the scFv in the offspring. In this study, we used scFv expression strategy by cloning an anti-CHIKV scFv into the NEGV genome. Using NEGV as a vehicle for paratransgensis is advantageous, because an isolate can infect multiple host species, and it is suspected to be vertically transmitted in mosquitoes and in theory could become established in multiple generations of the infected host species [16, 21, 47].

Modifications to the NEGV genome were tolerated as both cleaved and fusion proteins. Expression of extraneous proteins in viruses is common with 2A sequences to produce separate proteins or under a separate subgenomic promoter [48, 49]. However, extraneous proteins expressed as a fusion with a structural virus protein is uncommon. ORF3 is ∼25kDa and is suspected to be the membrane protein, the dominant structural protein; and ORF2 is ∼40kDa and is the putative glycoprotein predicted to form a bud projecting from one end of the virion [25]. The viability of the NEGV GFP- or scFv-fusion isolates was surprising because these inserts double the size of the membrane protein, which must interact with itself and ultimately support the projection of the glycoprotein. By using NEGV to express anti-CHIKV scFvs, the cleaved and fused inserts may provide distinct advantages. Cleaved scFvs are free to be transported around the cell, accessing many different locations where they may encounter CHIKV proteins. In contrast, fused proteins are bound to the membrane protein of NEGV and are limited to compartments of the cell where NEGV proteins are expressed and virions are assembled. In theory, increasing the concentration of the scFvs in specific areas of the cell, should inhibit CHIKV virion assembly and egression. By using both cleaved and fused NEGV isolates, the scFv sequence can also be easily replaced to target a new pathogen, adding to the versatility of this technique.

The current experiments demonstrate the ability of some negeviruses, both wild-type and paratransgenic isolates, to inhibit the replication in mosquito cells with co-infected arboviruses. The obvious next question is whether genetically altered negeviruses will survive and replicate in live mosquitoes; and if so, will they be vertically transmitted or transovarially transmitted in the insects? This will be our next area of investigation. If successful, then the use of paratransgenic negeviruses could be another novel method to alter the vector competence of mosquitoes for selected arboviruses.

## Acknowledgements

E.I.P. was supported by the Liverpool School of Tropical Medicine Director’s Catalyst Fund award. N.L.F. was supported by the NIH R01-AI095753-01A1 and R01-AI125902. G.L.H. was supported by the BBSRC (BB/T001240/1), the Royal Society Wolfson Fellowship (RSWF\R1\180013), NIH grants (R21AI138074 and R21AI129507), URKI (20197), and the NIHR (NIHR2000907).

## References

1. Calisher CH, Higgs S. The Discovery of Arthropod-Specific Viruses in Hematophagous Arthropods: An Open Door to Understanding the Mechanisms of Arbovirus and Arthropod Evolution? Annu Rev Entomol. 2018;63:87-103. Epub 2018/01/13. doi: 10.1146/annurev-ento-020117-043033. PubMed PMID: 29324047.

2. Ledermann JP, Suchman EL, Black WCt, Carlson JO. Infection and pathogenicity of the mosquito densoviruses AeDNV, HeDNV, and APeDNV in Aedes aegypti mosquitoes (Diptera: Culicidae). J Econ Entomol. 2004;97(6):1828-35. Epub 2005/01/26. doi: 10.1093/jee/97.6.1828. PubMed PMID: 15666733.

3. Carlson J, Suchman E, Buchatsky L. Densoviruses for control and genetic manipulation of mosquitoes. Adv Virus Res. 2006;68:361-92. Epub 2006/09/26. doi: 10.1016/S0065-3527(06)68010-X. PubMed PMID: 16997017.

4. Hirunkanokpun S, Carlson JO, Kittayapong P. Evaluation of mosquito densoviruses for controlling Aedes aegypti (Diptera: Culicidae): variation in efficiency due to virus strain and geographic origin of mosquitoes. Am J Trop Med Hyg. 2008;78(5):784-90. Epub 2008/05/07. PubMed PMID: 18458314.

5. Baidaliuk A, Miot EF, Lequime S, Moltini-Conclois I, Delaigue F, Dabo S, et al. Cell-Fusing Agent Virus Reduces Arbovirus Dissemination in Aedes aegypti Mosquitoes In Vivo. J Virol. 2019;93(18). Epub 2019/06/28. doi: 10.1128/JVI.00705-19. PubMed PMID: 31243123; PubMed Central PMCID: PMCPMC6714787.

6. Goenaga S, Kenney JL, Duggal NK, Delorey M, Ebel GD, Zhang B, et al. Potential for Co-Infection of a Mosquito-Specific Flavivirus, Nhumirim Virus, to Block West Nile Virus Transmission in Mosquitoes. Viruses. 2015;7(11):5801-12. Epub 2015/11/17. doi: 10.3390/v7112911. PubMed PMID: 26569286; PubMed Central PMCID: PMCPMC4664984.

7. Hall-Mendelin S, McLean BJ, Bielefeldt-Ohmann H, Hobson-Peters J, Hall RA, van den Hurk AF. The insect-specific Palm Creek virus modulates West Nile virus infection in and transmission by Australian mosquitoes. Parasit Vectors. 2016;9(1):414. Epub 2016/07/28. doi: 10.1186/s13071-016-1683-2. PubMed PMID: 27457250; PubMed Central PMCID: PMCPMC4960669.

8. Hobson-Peters J, Yam AW, Lu JW, Setoh YX, May FJ, Kurucz N, et al. A new insect-specific flavivirus from northern Australia suppresses replication of West Nile virus and Murray Valley encephalitis virus in co-infected mosquito cells. PLoS One. 2013;8(2):e56534. Epub 2013/03/06. doi: 10.1371/journal.pone.0056534. PubMed PMID: 23460804; PubMed Central PMCID: PMCPMC3584062.

9. Kenney JL, Solberg OD, Langevin SA, Brault AC. Characterization of a novel insect-specific flavivirus from Brazil: potential for inhibition of infection of arthropod cells with medically important flaviviruses. J Gen Virol. 2014;95(Pt 12):2796-808. Epub 2014/08/26. doi: 10.1099/vir.0.068031-0. PubMed PMID: 25146007; PubMed Central PMCID: PMCPMC4582674.

10. Romo H, Kenney JL, Blitvich BJ, Brault AC. Restriction of Zika virus infection and transmission in Aedes aegypti mediated by an insect-specific flavivirus. Emerg Microbes Infect. 2018;7(1):181. Epub 2018/11/16. doi: 10.1038/s41426-018-0180-4. PubMed PMID: 30429457; PubMed Central PMCID: PMCPMC6235874.

11. Nasar F, Erasmus JH, Haddow AD, Tesh RB, Weaver SC. Eilat virus induces both homologous and heterologous interference. Virology. 2015;484:51-8. Epub 2015/06/13. doi: 10.1016/j.virol.2015.05.009. PubMed PMID: 26068885; PubMed Central PMCID: PMCPMC4567418.

12. Burivong P, Pattanakitsakul SN, Thongrungkiat S, Malasit P, Flegel TW. Markedly reduced severity of Dengue virus infection in mosquito cell cultures persistently infected with Aedes albopictus densovirus (AalDNV). Virology. 2004;329(2):261-9. Epub 2004/11/03. doi: 10.1016/j.virol.2004.08.032. PubMed PMID: 15518806.

13. Schultz MJ, Frydman HM, Connor JH. Dual Insect specific virus infection limits Arbovirus replication in Aedes mosquito cells. Virology. 2018;518:406-13. Epub 2018/04/07. doi: 10.1016/j.virol.2018.03.022. PubMed PMID: 29625404.

14. Zhang G, Asad S, Khromykh AA, Asgari S. Cell fusing agent virus and dengue virus mutually interact in Aedes aegypti cell lines. Sci Rep. 2017;7(1):6935. Epub 2017/08/02. doi: 10.1038/s41598-017-07279-5. PubMed PMID: 28761113; PubMed Central PMCID: PMCPMC5537255.

15. Bolling BG, Olea-Popelka FJ, Eisen L, Moore CG, Blair CD. Transmission dynamics of an insect-specific flavivirus in a naturally infected Culex pipiens laboratory colony and effects of co-infection on vector competence for West Nile virus. Virology. 2012;427(2):90-7. Epub 2012/03/20. doi: 10.1016/j.virol.2012.02.016. PubMed PMID: 22425062; PubMed Central PMCID: PMCPMC3329802.

16. Vasilakis N, Forrester NL, Palacios G, Nasar F, Savji N, Rossi SL, et al. Negevirus: a proposed new taxon of insect-specific viruses with wide geographic distribution. J Virol. 2013;87(5):2475-88. Epub 2012/12/21. doi: 10.1128/JVI.00776-12. PubMed PMID: 23255793; PubMed Central PMCID: PMCPMC3571365.

17. Carapeta S, do Bem B, McGuinness J, Esteves A, Abecasis A, Lopes A, et al. Negeviruses found in multiple species of mosquitoes from southern Portugal: Isolation, genetic diversity, and replication in insect cell culture. Virology. 2015;483:318-28. Epub 2015/06/10. doi: 10.1016/j.virol.2015.04.021. PubMed PMID: 26057025.

18. Charles J, Tangudu CS, Hurt SL, Tumescheit C, Firth AE, Garcia-Rejon JE, et al. Detection of novel and recognized RNA viruses in mosquitoes from the Yucatan Peninsula of Mexico using metagenomics and characterization of their in vitro host ranges. J Gen Virol. 2018;99(12):1729-38. Epub 2018/11/10. doi: 10.1099/jgv.0.001165. PubMed PMID: 30412047; PubMed Central PMCID: PMCPMC6537628.

19. Kondo H, Chiba S, Maruyama K, Andika IB, Suzuki N. A novel insect-infecting virga/nege-like virus group and its pervasive endogenization into insect genomes. Virus Res. 2019;262:37-47. Epub 2017/11/25. doi: 10.1016/j.virusres.2017.11.020. PubMed PMID: 29169832.

20. Kondo H, Fujita M, Hisano H, Hyodo K, Andika IB, Suzuki N. Virome Analysis of Aphid Populations That Infest the Barley Field: The Discovery of Two Novel Groups of Nege/Kita-Like Viruses and Other Novel RNA Viruses. Front Microbiol. 2020;11:509. Epub 2020/04/23. doi: 10.3389/fmicb.2020.00509. PubMed PMID: 32318034; PubMed Central PMCID: PMCPMC7154061.

21. Nunes MRT, Contreras-Gutierrez MA, Guzman H, Martins LC, Barbirato MF, Savit C, et al. Genetic characterization, molecular epidemiology, and phylogenetic relationships of insect-specific viruses in the taxon Negevirus. Virology. 2017;504:152-67. Epub 2017/02/15. doi: 10.1016/j.virol.2017.01.022. PubMed PMID: 28193550; PubMed Central PMCID: PMCPMC5394984.

22. De MirandaJRH, H.; Onorati, P.; Stephan, J.; Karlberg, O.; Bylund, H.; Terenius, O. Characterization of a Novel RNA Virus Discovered in the Autumnal Moth Epirrita autumnata in Sweden. Viruses. 2017;9. doi: https://doi.org/10.3390/v9080214.

23. Kallies R, Kopp A, Zirkel F, Estrada A, Gillespie TR, Drosten C, et al. Genetic characterization of goutanap virus, a novel virus related to negeviruses, cileviruses and higreviruses. Viruses. 2014;6(11):4346-57. Epub 2014/11/15. doi: 10.3390/v6114346. PubMed PMID: 25398046; PubMed Central PMCID: PMCPMC4246226.

24. Lu G, Ye ZX, He YJ, Zhang Y, Wang X, Huang HJ, et al. Discovery of Two Novel Negeviruses in a Dungfly Collected from the Arctic. Viruses. 2020;12(7). Epub 2020/07/02. doi: 10.3390/v12070692. PubMed PMID: 32604989.

25. Colmant AMG, O’Brien CA, Newton ND, Watterson D, Hardy J, Coulibaly F, et al. Novel monoclonal antibodies against Australian strains of negeviruses and insights into virus structure, replication and host -restriction. J Gen Virol. 2020;101(4):440-52. Epub 2020/02/01. doi: 10.1099/jgv.0.001388. PubMed PMID: 32003709.

26. Nabeshima T, Inoue S, Okamoto K, Posadas-Herrera G, Yu F, Uchida L, et al. Tanay virus, a new species of virus isolated from mosquitoes in the Philippines. J Gen Virol. 2014;95(Pt 6):1390-5. Epub 2014/03/22. doi: 10.1099/vir.0.061887-0. PubMed PMID: 24646751.

27. O’Brien CA, McLean BJ, Colmant AMG, Harrison JJ, Hall-Mendelin S, van den Hurk AF, et al. Discovery and Characterisation of Castlerea Virus, a New Species of Negevirus Isolated in Australia. Evol Bioinform Online. 2017;13:1176934317691269. Epub 2017/05/05. doi: 10.1177/1176934317691269. PubMed PMID: 28469377; PubMed Central PMCID: PMCPMC5395271.

28. Shi M, Neville P, Nicholson J, Eden JS, Imrie A, Holmes EC. High-Resolution Metatranscriptomics Reveals the Ecological Dynamics of Mosquito-Associated RNA Viruses in Western Australia. J Virol. 2017;91(17). Epub 2017/06/24. doi: 10.1128/JVI.00680-17. PubMed PMID: 28637756; PubMed Central PMCID: PMCPMC5553174.

29. Atoni E, Wang Y, Karungu S, Waruhiu C, Zohaib A, Obanda V, et al. Metagenomic Virome Analysis of Culex Mosquitoes from Kenya and China. Viruses. 2018;10(1). Epub 2018/01/13. doi: 10.3390/v10010030. PubMed PMID: 29329230; PubMed Central PMCID: PMCPMC5795443.

30. Walker T, Jeffries CL, Mansfield KL, Johnson N. Mosquito cell lines: history, isolation, availability and application to assess the threat of arboviral transmission in the United Kingdom. Parasit Vectors. 2014;7:382. Epub 2014/08/22. doi: 10.1186/1756-3305-7-382. PubMed PMID: 25141888; PubMed Central PMCID: PMCPMC4150944.

31. Gorchakov RV, Tesh RB, Weaver SC, Nasar F. Generation of an infectious Negev virus cDNA clone. J Gen Virol. 2014;95(Pt 9):2071-4. Epub 2014/06/01. doi: 10.1099/vir.0.066019-0. PubMed PMID: 24878640; PubMed Central PMCID: PMCPMC4135092.

32. Levitt NH, Ramsburg HH, Hasty SE, Repik PM, Cole FE, Jr., Lupton HW. Development of an attenuated strain of chikungunya virus for use in vaccine production. Vaccine. 1986;4(3):157-62. Epub 1986/09/01. doi: 10.1016/0264-410x(86)90003-4. PubMed PMID: 3020820.

33. Smith DR, Adams AP, Kenney JL, Wang E, Weaver SC. Venezuelan equine encephalitis virus in the mosquito vector Aedes taeniorhynchus: infection initiated by a small number of susceptible epithelial cells and a population bottleneck. Virology. 2008;372(1):176-86. Epub 2007/11/21. doi: 10.1016/j.virol.2007.10.011. PubMed PMID: 18023837; PubMed Central PMCID: PMCPMC2291444.

34. Kinney RM, Chang GJ, Tsuchiya KR, Sneider JM, Roehrig JT, Woodward TM, et al. Attenuation of Venezuelan equine encephalitis virus strain TC-83 is encoded by the 5’-noncoding region and the E2 envelope glycoprotein. J Virol. 1993;67(3):1269-77. Epub 1993/03/01. doi: 10.1128/JVI.67.3.1269-1277.1993. PubMed PMID: 7679745; PubMed Central PMCID: PMCPMC237493.

35. Fox JM, Long F, Edeling MA, Lin H, van Duijl-Richter MKS, Fong RH, et al. Broadly Neutralizing Alphavirus Antibodies Bind an Epitope on E2 and Inhibit Entry and Egress. Cell. 2015;163(5):1095-107. Epub 2015/11/11. doi: 10.1016/j.cell.2015.10.050. PubMed PMID: 26553503; PubMed Central PMCID: PMCPMC4659373.

36. Cross ST, Kapuscinski ML, Perino J, Maertens BL, Weger-Lucarelli J, Ebel GD, et al. Co-Infection Patterns in Individual Ixodes scapularis Ticks Reveal Associations between Viral, Eukaryotic and Bacterial Microorganisms. Viruses. 2018;10(7). Epub 2018/07/25. doi: 10.3390/v10070388. PubMed PMID: 30037148; PubMed Central PMCID: PMCPMC6071216.

37. Hegde S, Khanipov K, Albayrak L, Golovko G, Pimenova M, Saldana MA, et al. Microbiome Interaction Networks and Community Structure From Laboratory-Reared and Field-Collected Aedes aegypti, Aedes albopictus, and Culex quinquefasciatus Mosquito Vectors. Front Microbiol. 2018;9:2160. Epub 2018/09/27. doi: 10.3389/fmicb.2018.02160. PubMed PMID: 30250462; PubMed Central PMCID: PMCPMC6140713.

38. Nanfack-Minkeu F, Mitri C, Bischoff E, Belda E, Casademont I, Vernick KD. Interaction of RNA viruses of the natural virome with the African malaria vector, Anopheles coluzzii. Sci Rep. 2019;9(1):6319. Epub 2019/04/21. doi: 10.1038/s41598-019-42825-3. PubMed PMID: 31004099; PubMed Central PMCID: PMCPMC6474895.

39. Kuwata R, Isawa H, Hoshino K, Sasaki T, Kobayashi M, Maeda K, et al. Analysis of Mosquito-Borne Flavivirus Superinfection in Culex tritaeniorhynchus (Diptera: Culicidae) Cells Persistently Infected with Culex Flavivirus (Flaviviridae). J Med Entomol. 2015;52(2):222-9. Epub 2015/09/04. doi: 10.1093/jme/tju059. PubMed PMID: 26336307.

40. Newman CM, Krebs BL, Anderson TK, Hamer GL, Ruiz MO, Brawn JD, et al. Culex Flavivirus During West Nile Virus Epidemic and Interepidemic Years in Chicago, United States. Vector Borne Zoonotic Dis. 2017;17(8):567-75. Epub 2017/06/20. doi: 10.1089/vbz.2017.2124. PubMed PMID: 28628366; PubMed Central PMCID: PMCPMC5564057.

41. Talavera S, Birnberg L, Nunez AI, Munoz-Munoz F, Vazquez A, Busquets N. Culex flavivirus infection in a Culex pipiens mosquito colony and its effects on vector competence for Rift Valley fever phlebovirus. Parasit Vectors. 2018;11(1):310. Epub 2018/05/25. doi: 10.1186/s13071-018-2887-4. PubMed PMID: 29792223; PubMed Central PMCID: PMCPMC5966921.

42. Kent RJ, Crabtree MB, Miller BR. Transmission of West Nile virus by Culex quinquefasciatus say infected with Culex Flavivirus Izabal. PLoS Negl Trop Dis. 2010;4(5):e671. Epub 2010/05/11. doi: 10.1371/journal.pntd.0000671. PubMed PMID: 20454569; PubMed Central PMCID: PMCPMC2864301.

43. Patterson EI, Villinger J, Muthoni JN, Dobel-Ober L, Hughes GL. Exploiting insect-specific viruses as a novel strategy to control vector-borne disease. Curr Opin Insect Sci. 2020;39:50-6. Epub 2020/04/12. doi: 10.1016/j.cois.2020.02.005. PubMed PMID: 32278312; PubMed Central PMCID: PMCPMC7302987.

44. Jiggins FM. The spread of Wolbachia through mosquito populations. PLoS Biol. 2017;15(6):e2002780. Epub 2017/06/02. doi: 10.1371/journal.pbio.2002780. PubMed PMID: 28570608; PubMed Central PMCID: PMCPMC5453404.

45. Gantz VM, Jasinskiene N, Tatarenkova O, Fazekas A, Macias VM, Bier E, et al. Highly efficient Cas9-mediated gene drive for population modification of the malaria vector mosquito Anopheles stephensi. Proc Natl Acad Sci U S A. 2015;112(49):E6736-43. Epub 2015/11/26. doi: 10.1073/pnas.1521077112. PubMed PMID: 26598698; PubMed Central PMCID: PMCPMC4679060.

46. Buchman A, Gamez S, Li M, Antoshechkin I, Li HH, Wang HW, et al. Broad dengue neutralization in mosquitoes expressing an engineered antibody. PLoS Pathog. 2020;16(1):e1008103. Epub 2020/01/17. doi: 10.1371/journal.ppat.1008103. PubMed PMID: 31945137; PubMed Central PMCID: PMCPMC6964813 J.E.C. has served as a consultant for Takeda Vaccines, Sanofi Pasteur, Pfizer, and Novavax, is on the Scientific Advisory Boards of CompuVax, GigaGen, Meissa Vaccines, and is the Founder of IDBiologics, Inc. All other authors declare no competing financial interests.

47. Kawakami K, Kurnia YW, Fujita R, Ito T, Isawa H, Asano S, et al. Characterization of a novel negevirus isolated from Aedes larvae collected in a subarctic region of Japan. Arch Virol. 2016;161(4):801-9. Epub 2015/12/22. doi: 10.1007/s00705-015-2711-9. PubMed PMID: 26687585.

48. Kautz TF, Jaworski E, Routh A, Forrester NL. A Low Fidelity Virus Shows Increased Recombination during the Removal of an Alphavirus Reporter Gene. Viruses. 2020;12(6). Epub 2020/06/25. doi: 10.3390/v12060660. PubMed PMID: 32575413; PubMed Central PMCID: PMCPMC7354468.

49. Suzuki Y, Niu G, Hughes GL, Rasgon JL. A viral over-expression system for the major malaria mosquito Anopheles gambiae. Sci Rep. 2014;4:5127. Epub 2014/05/31. doi: 10.1038/srep05127. PubMed PMID: 24875042; PubMed Central PMCID: PMCPMC4038844.

